# Enhanced preference for delayed rewards requires the basolateral amygdala and retrosplenial cortex

**DOI:** 10.1101/2021.01.14.426754

**Authors:** Merridee J. Lefner, Alexa P. Magnon, James M. Gutierrez, Matthew R. Lopez, Matthew J. Wanat

## Abstract

‘Sunk’ or irrecoverable costs imposed on a subject can impact reward-based decisions. However, it is not known if these incurred costs elicit a sustained change in reward value. To address this, we examined if sunk temporal costs subsequently alter reward consumption and reward preference in rats. We first identified the relative preference between different flavored food pellets during a free-feeding consumption test. Animals were then trained to experience the initial less preferred reward after long delays and the initial preferred reward after short delays. This training regimen enhanced the consumption and preference for the initial less desirable food reward. We probed whether this change in reward preference involved neural systems that contribute to reward valuation. Pharmacological manipulations and site-specific lesions were performed to examine the potential involvement of the dopamine system, the orbitofrontal cortex (OFC), the basolateral amygdala (BLA), and the retrosplenial cortex (RSC). The change in reward preference was unaffected by systemic dopamine receptor antagonism or OFC lesions. In contrast, lesions of the BLA or the RSC prevented the enhanced consumption and preference for the initial less desirable reward. These findings demonstrate that both the BLA and RSC participate in how sunk temporal costs alter reward value and reward preference.

**Significance Statement:** From an economic standpoint, only future costs should factor into one’s decisions. However, behavioral evidence across species illustrates that *past* costs can alter decisions. The goal of this study was to identify the neural systems responsible for past costs influencing subsequent actions. We demonstrate that delivering an initially less desirable reward after long delays (high temporal costs) subsequently increases the consumption and preference for that reward. Furthermore, we identified the basolateral amygdala and the retrosplenial cortex as essential nuclei for mediating change in reward preference elicited by past temporal costs.

## Introduction

Reward-based decisions are driven by cost-benefit analyses^1,2^. A purely economic decision should be influenced solely by the future costs associated with earning a reward^3,4^. However, behavioral evidence across species demonstrate that *past* costs influence decision-making policies^5–12^. These ‘sunk costs’ are responsible for the enhanced preference for conditioned stimuli associated with rewards that follow substantial effort or delays^5–7^. Furthermore, sunk costs can result in fewer rewards earned, as subjects will persist in an action even though it is advantageous to disengage from their current behavior^11,12^. The effect of sunk costs on behavior has been primarily studied in tasks that involve opportunity costs, in which choosing one course of action comes at the expense of an alternative outcome^11^. However, this approach cannot determine if sunk costs increase the value of the chosen outcome and/or decrease the value of the alternative outcome.

To address this, we developed a rodent task to determine how sunk temporal costs prior to a reward delivery subsequently alters reward consumption and preference. We first identified the relative preference between different flavored food rewards in a free-feeding test. Rats then underwent training sessions in which the initial less preferred reward was delivered after long delays (high temporal cost), while the initial preferred reward was delivered after short delays (low temporal cost). The impact of these training sessions on food consumption and reward preference was then assessed in a second free-feeding test performed one day after the last training session. In this manner, we could test the hypothesis that high temporal costs imposed during training sessions subsequently increases the value (consumption) and relative preference for the initial less desirable reward.

By utilizing this behavioral task, we could identify brain regions responsible for changes in reward preference elicited by sunk temporal costs. Neural systems involved with reward valuation and learning are potential candidates for mediating changes in reward preference. These neurobiological processes are evident throughout the brain. For example, dopamine neurons convey the relative value between reward options and contribute to appetitive learning^13–18^. Pyramidal neurons in the orbitofrontal cortex (OFC) encode value-based parameters and participate in reward-learning and decision-making^19–25^. The basolateral amygdala (BLA) is also a candidate due to its role in learning, reward valuation, and reward seeking^26–31^. Finally, the retrosplenial cortex (RSC) could be required for long-lasting changes in reward preference since RSC neurons contribute to memory formation and encode value-related signals^32–35^. Therefore, in this study we performed pharmacological manipulations and site-specific lesions to establish whether changes in preference elicited by incurred temporal costs require dopamine signaling, the OFC, the BLA, and/or the RSC.

## Methods

### Subjects and surgery

All procedures were approved by the Institutional Animal Care and Use Committee at the University of Texas at San Antonio. 300-350 g male and female Sprague-Dawley rats (Charles River, MA) were pair-housed upon arrival and given ad libitum access to water and chow and maintained on a 12-hour light/dark cycle.

All surgeries were performed under isoflurane anesthesia and drug infusions were delivered at a rate of 0.1μL/min. Surgical coordinates and injection volumes were based on prior research^25,31,36^. OFC-lesioned rats received injections of NMDA (12.5 μg/μL in saline vehicle; Tocris) at the following locations (relative to bregma): 3.0 mm AP, ± 3.2 mm ML, −5.2 mm DV (0.05 μL); 3.0 mm AP, ± 4.2 mm ML, −5.2 mm DV (0.1 μL); 4.0 mm AP, ± 2.2 mm ML, −3.8 mm DV (0.1 μL); 4.0 mm AP, ± 3.7 mm ML, −3.8 mm DV (0.1 μL). BLA-lesioned rats received injections of NMDA (20 μg/μL) at the following locations: −3.3 mm AP, ± 4.6 mm ML, −8.6 mm DV (0.2 μL); −3.3 mm AP, ± 4.6 mm ML, −8.4 mm DV (0.1 μL). RSC-lesioned rats received injections of NMDA (20 μg/μL) at the following locations: −1.6 mm AP, ± 0.5 mm ML, −1.3 mm DV (0.26 μL); −2.8 mm AP, ± 0.5 mm ML, −1.3 mm DV (0.26 μL); −4.0 mm AP, ± 0.5 mm ML, −1.3 mm DV (0.26 μL injection); −5.3 mm AP, ± 0.5 mm ML, −2.0 mm DV (0.26 μL injection). All sham surgeries involved lowering the injector to the respective injection sites^25,31,36^. Animals recovered for >1 week following surgery before beginning training.

### Training

Rats were placed and maintained on mild food restriction (~15 g/day of standard lab chow) to target 90% free-feeding weight, allowing for an increase of 1.5% per week. Behavioral sessions were performed in chambers that had grid floors, a house light, and two food trays on a single wall. In free-feeding sessions, plastic barriers were placed over the food trays. Additionally, a plastic insert was placed over the grid floors that contained two fixed cups in which the food pellets were placed. Experimental 45 mg sucrose pellets (chocolate flavor #F0025 and banana flavor #F0024; Bio-Serv) were placed in their home cages to minimize neophobia. Rats first underwent a free-feeding session (10 mins) in which a single food pellet flavor was offered (6.5 g total). On the following day, rats underwent a second free-feeding session in which the alternate flavor was offered (ordering counterbalanced between animals). For the free-feeding preference test, rats were allowed 10 min to consume both chocolate and banana food pellets that were freely available in cups affixed to the floor. To ensure an ample supply of food, we provided 13 g of each flavor, which was 3 g higher than the maximal amount consumed in pilot studies. We identified which reward flavor was the Initial Preferred and the Initial Less Preferred based on the food consumed during this preference test.

Rats next underwent training sessions (1/day) in which one of the rewards was delivered non-contingently for a total of 50 pellets per session. The Initial Less Preferred reward was delivered after a 60 ±5 s inter-trial interval (ITI) in Long Delay sessions. The Initial Preferred reward was delivered after a 30 ±5 s ITI in Short Delay sessions. There were a total of 10 training sessions, which alternated between Long and Short Delay sessions with the first session counterbalanced between animals. Rats underwent a second free-feeding preference test following this training regimen (**Fig. 1A**). In a control experiment, rats were trained as described above except that a 45 ±5 s ITI was utilized for both the Initial Less Preferred and the Initial Preferred reward training sessions (see **Fig. 2A**). Injections of flupenthixol (225 μg/kg i.p., Tocris) or saline were administered 1 hour prior to training sessions, based on established research^37^.

**Figure 1.**
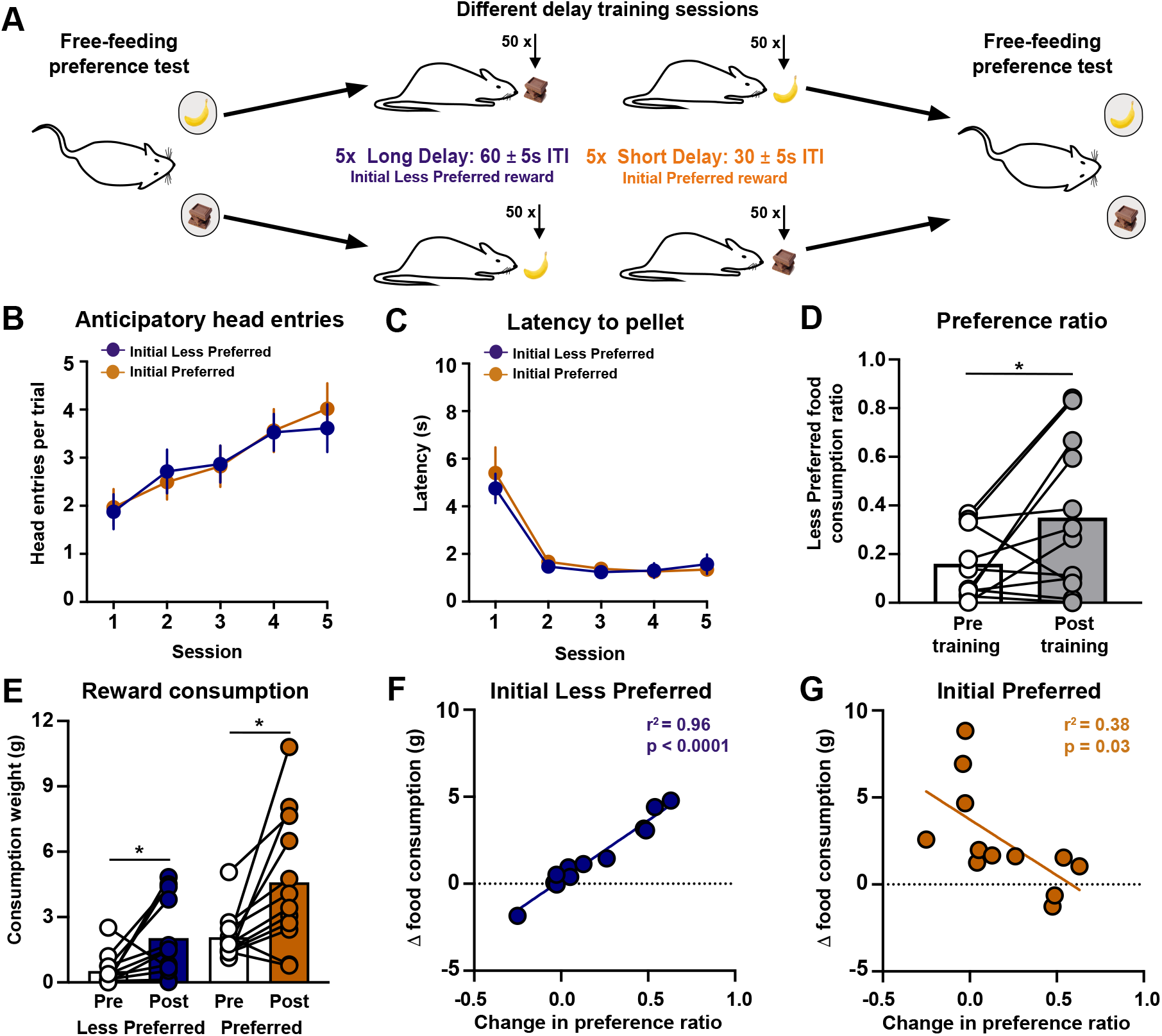
Increased preference for rewards delivered after high temporal costs. (A) Training schematic. (B) Anticipatory head entries into the food port during the 5s prior to reward delivery for the Initial Less Preferred (Long Delay) and Initial Preferred (Short Delay) training sessions. (C) Latency to make a head entry into the food port after a reward is delivered for the Initial Less Preferred and Initial Preferred training sessions. (D) Preference ratio plotted as the amount of the Initial Less Preferred food consumed out of the total food consumed during preference tests. (E) Reward consumption for each flavor during the preference tests. (F,G) Linear regression relating the change in food consumption as a function of the change in the preference ratio. * *p*< 0.05.

**Figure 2.**
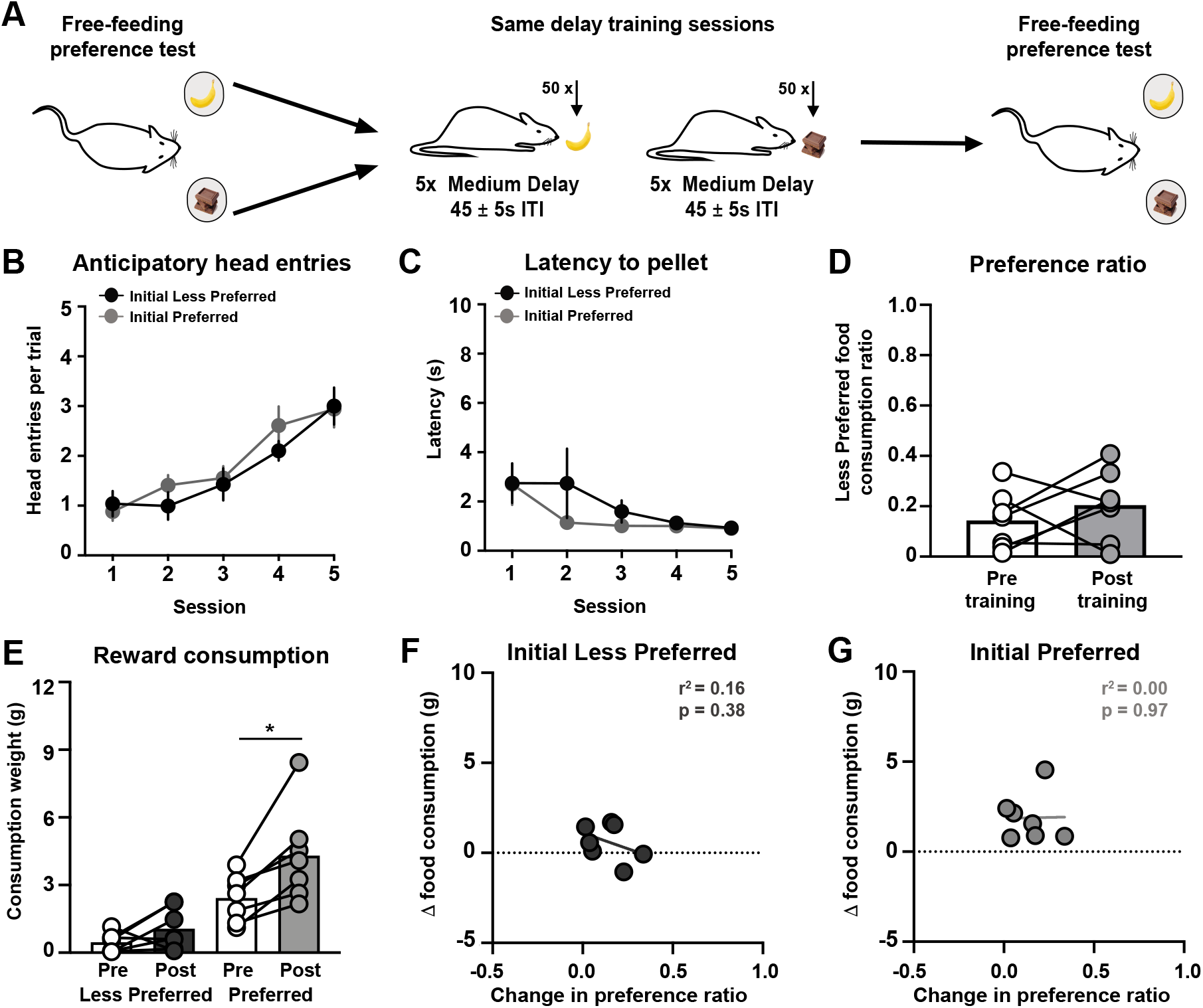
No change in reward preference following same delay training sessions. (A) Training schematic. (B) Anticipatory head entries into the food port during the 5s prior to reward delivery for the Initial Less Preferred (Medium Delay) and Initial Preferred (Medium Delay) training sessions. (C) Latency to make a head entry into food port after a reward is delivered for the Initial Less Preferred and the Initial Preferred training sessions. (D) Preference ratio. (E) Reward consumption for each flavor during the preference tests. (F,G) Linear regression relating the change in food consumption as a function of the change in the preference ratio. * p < 0.05.

### Data analysis

The latency to respond was calculated as the length of time to make a head entry into the food port after a pellet was delivered. The latency measurement was capped at 25 s (shortest possible trial duration) to account for the rare occasion when a pellet was not retrieved before the subsequent trial. Anticipatory head entries were measured as the number of head entries into the food port during the 5 s preceding the reward delivery. The preference ratio was calculated as the amount consumed of the Initial Less Preferred reward relative to the total food consumed during the free-feeding test. We performed statistical analyses using GraphPad Prism 8. The effect of training on behavioral outcomes were analyzed using a paired t-test or a mixed-effects model fit (restricted maximum likelihood method), repeated measures where appropriate, followed by a post-hoc Sidak’s test. The Geisser-Greenhouse correction was applied to address unequal variances between groups. The significance level was set to α = 0.05 for all tests. The full list of statistical analyses is in **Supplementary Data**.

### Histology

Rats were intracardially perfused with 4% paraformaldehyde, and brains were removed and postfixed for at least 24 hours. Brains were subsequently placed in 15% and 30% sucrose solutions in phosphate-buffered saline. Brains were then flash frozen on dry ice, coronally sectioned, and stained with cresyl violet to verify the location and spread of the surgical lesions.

## Results

We developed a rodent task to examine how sunk temporal costs impact reward value and preference. Rats first underwent a free-feeding preference test between food rewards that had an identical nutritional profile but differed in flavor (chocolate or banana). We identified which was the Initial Preferred and the Initial Less Preferred flavor for each rat based on the food consumed during this free-feeding test. Rats then underwent training sessions in which one of the food rewards was delivered non-contingently for a total of 50 pellets per session. The Initial Less Preferred reward was delivered after a Long Delay (60 ±5 s ITI), while the Initial Preferred reward was delivered after a Short Delay (30 ±5 s ITI) in separate sessions. Rats underwent a total of 10 training sessions, which alternated between Long and Short Delay sessions, with the first session counterbalanced between animals. To assess the impact of this training regimen on food consumption and reward preference, rats underwent a second free-feeding preference test following training (different delay training; **Fig. 1A**).

Food pellets were delivered non-contingently and were not preceded by reward-predictive cues during training sessions. Across sessions, rats increased anticipatory head entries into the food port during the 5 s prior to the reward delivery, but there was no difference between Long and Short Delay sessions (two-way mixed-effects analysis; session effect: *F*_(1.894, 20.83)_ = 6.52, *p* = 0.007; delay effect: *F*_(1, 11)_ = 0.05, *p* = 0.8337; interaction effect: *F*_(2.661, 29.27)_ = 0.92, *p* = 0.4334; **Fig. 1B**). Rats also decreased the post-reward latency into the food port across training, with no difference between session type (two-way mixed-effects analysis; session effect: *F*_(1.316, 14.48)_ = 54.17, *p* < 0.0001; delay effect: *F*_(1, 11)_ = 0.16, *p* = 0.7002; interaction effect: *F*_(1.30, 12.43)_ = 0.23, *p* = 0.6672; n = 12 rats; **Fig. 1C**). Even though there were no behavioral differences during Long and Short Delay sessions, this training regimen increased the rats’ preference for the Initial Less Preferred reward (paired t-test: *t*_(11)_ = 2.33, *p* = 0.0402; **Fig. 1D**) due to increased consumption of that reward option (post-hoc Sidak’s test: *t*_(11)_ = 2.65, *p* = 0.0451; **Fig. 1E**). This cohort also exhibited an increased consumption of the Initial Preferred reward after training (post-hoc Sidak’s test: *t*_(11)_ = 2.98, *p* = 0.0251; **Fig. 1E**). We next examined how the change in reward preference correlated with the change in food consumption across subjects. A stronger bias toward the Initial Less Preferred reward was positively related to the consumption of that reward option (r^2^ = 0.96, *p* < 0.0001, **Fig. 1F**) and inversely related to the consumption of the Initial Preferred reward (r^2^ = 0.38, *p* = 0.03, **Fig. 1G**).

The change in reward preference following the different delay training regimen could be due to the temporal costs imposed during training sessions or alternatively due to increased exposure with the Initial Less Preferred reward. To address this possibility, we trained a separate group of rats as described above except that a 45 ± 5 s ITI was used for both the Initial Less Preferred and Initial Preferred reward training sessions (same delay training; **Fig. 2A**). Rats increased anticipatory head entries into the food port across sessions, with no difference between session type (two-way mixed-effects analysis; session effect: *F*_(2.831, 16.99)_ = 37.89, *p* < 0.0001; delay effect: *F*_(1, 6)_ = 1.20, *p* = 0.3146; interaction effect: *F*_(2.297, 13.78)_ = 1.58, *p* = 0.2414; **Fig. 2B**). There was no difference in the latency between sessions (two-way mixed-effects analysis; session effect:*F*_(1.341, 8.049)_ = 2.97, *p* = 0.1178; delay effect: *F*_(1, 6)_ = 14.93, *p* = 0.2675; interaction effect: *F*_(2.094, 12.04)_ = 0.62, *p* = 0.5636; n = 7 rats; **Fig. 2C**). Rats undergoing the same delay training regimen did not alter their initial reward preference (paired t-test: *t*_(6)_ = 0.91, *p* = 0.40; **Fig. 2D**), as there was no change in consumption of the Initial Less Preferred reward (post-hoc Sidak’s test: *t*_(6)_ = 1.56, *p* = 0.3104; **Fig. 2E**) and an increased consumption of the Initial Preferred reward (post-hoc Sidak’s test: *t*_(6)_ = 3.71*, p* = 0.0199; **Fig. 2E**). Finally, there was no relationship between changes in preference and the consumption of the Initial Less Preferred reward (r^2^ = 0.16, *p* = 0.38, **Fig. 2F**) or the Initial Preferred reward (r^2^ = 0.00, *p* = 0.97, **Fig. 2G**) when the temporal cost was held constant across all training sessions. These experiments collectively demonstrate that the enhanced preference for the Initial Less Preferred reward following the different delay training regimen was due to differential temporal costs experienced during training sessions and not a result of increased exposure to the rewards over training.

We next sought to identify the neural systems responsible for sunk temporal costs altering reward preference. Dopamine neurons respond to rewards to convey the relative preference between options as well as changes in reward value^13–16,38^. Additionally, the dopamine system contributes to reward learning^17,18^. Therefore, the increased preference for rewards that follow greater delays could be mediated by dopamine signaling. To address this possibility, we systemically administered flupenthixol prior to the different delay training sessions and examined the consequence on reward preference (**Fig. 3A**). Anticipatory head entries into the food port were disrupted by flupenthixol treatment across sessions (three-way mixed-effects analysis; session effect: *F*_(3.020, 66.43)_ = 22.26, *p* < 0.0001; treatment effect: *F*_(1, 87)_ = 11.22, *p* = 0.0012; delay effect: *F*_(1, 22)_ = 0.05, *p* = 0.8188; session x treatment effect: *F*_(4, 87)_ = 11.22, *p* < 0.0001; interaction effect: *F*_(4, 87)_ = 0.32, *p* = 0.8644; **Fig. 3B**). Flupenthixol treatment also slowed the time to retrieve the reward during training sessions (three-way mixed-effects analysis; session effect: *F*_(4, 88)_ = 43.04, *p* < 0.0001; treatment effect: *F*_(1, 22)_ = 6.18, *p* = 0.016; delay effect: *F*_(1, 22)_ = 0.02, *p* = 0.8840; interaction effect: *F*_(4, 88)_ = 1.53, *p* = 0.2010; n = 12 saline, n = 12 flupenthixol rats; **Fig. 3C**). However, antagonizing dopamine receptors during training sessions did not prevent the increased preference for the Initial Less Preferred reward (two-way mixed-effects analysis; training effect: *F*_(1, 22)_ = 30.86, *p* < 0.0001; treatment effect: *F*_(1, 22)_ = 0.11, *p* = 0.7384; interaction effect: *F*_(1, 22)_ = 0.77, *p* = 0.3901; post-hoc Sidak’s test; saline: *t*_(22)_ = 4.55, *p* = 0.0003; flupenthixol: *t*_(22)_ = 3.31, *p* = 0.0064; **Fig. 3D**). Both saline- and flupenthixol-treated rats increased the consumption of the Initial Less Preferred reward (post-hoc Sidak’s test; saline: *t*_(11)_ = 4.92, *p* = 0.0009; flupenthixol: *t*_(11)_ = 3.54, *p* = 0.0092; **Fig. 3E**) with no change in consumption of the Initial Preferred reward (post-hoc Sidak’s test; saline: *t*_(11)_ = 0.91*, p* = 0.6169; flupenthixol: *t*_(11)_ = 1.77, *p* = 0.1984; **Fig. 3E**). The change in reward preference was positively related to the consumption of the Initial Less Preferred reward (saline: r^2^ = 0.72, *p* < 0.001, flupenthixol: r^2^ = 0.74, *p* < 0.001; **Fig. 3F**) and inversely related to the consumption of the Initial Preferred reward in both saline- and flupenthixol-treated animals (saline: r^2^ = 0.54, *p* < 0.01, flupenthixol: r^2^ = 0.46, *p* = 0.02, **Fig. 3G**). These data collectively demonstrate that the dopamine system regulates both anticipatory head entries and the latency to respond during training sessions, but does not mediate the change in reward preference elicited by sunk temporal costs.

**Figure 3.**
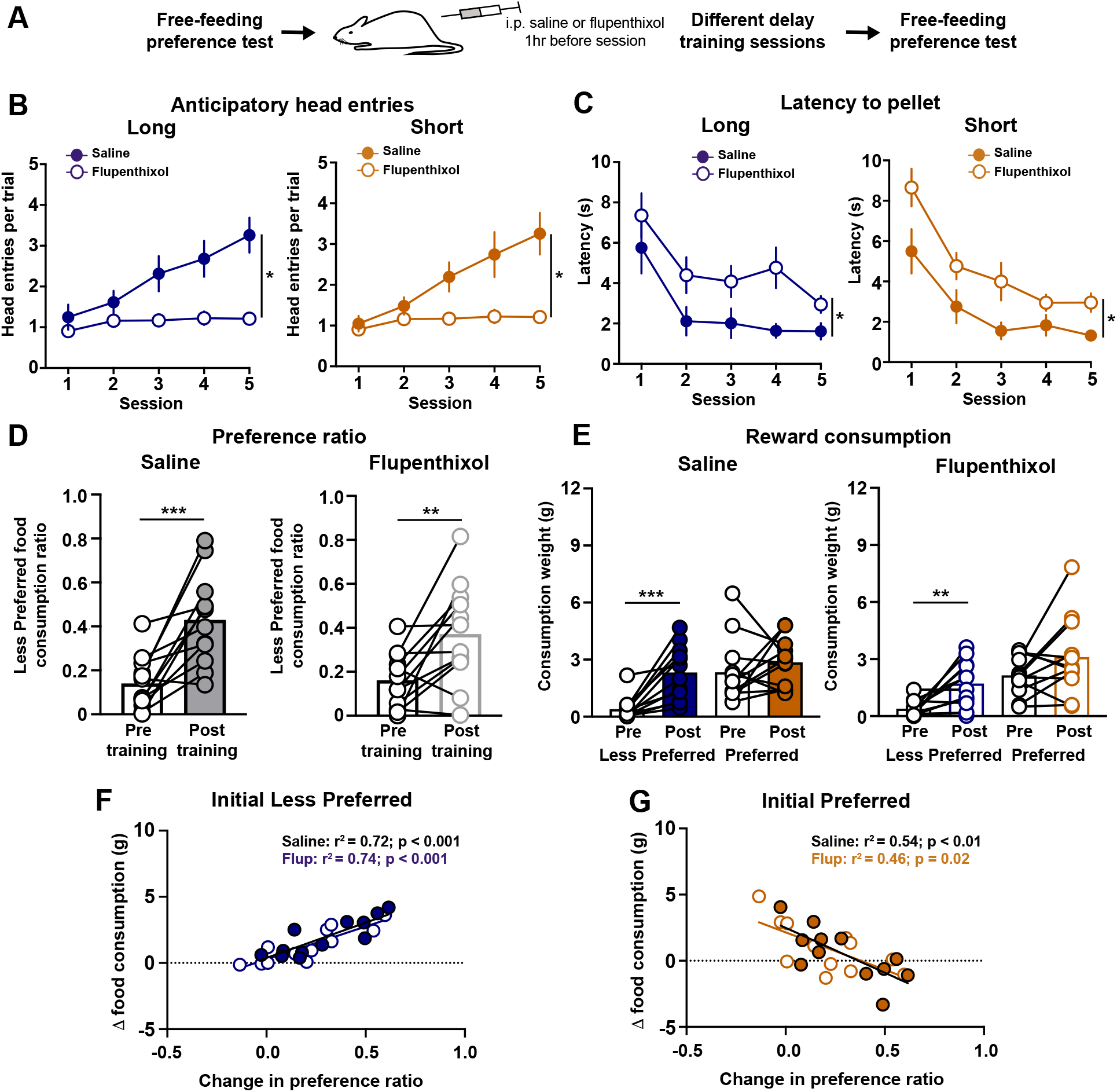
The change in reward preference does not involve dopamine signaling. (A) Training schematic. (B) Latency to make a head entry into the food port after a reward is delivered during training sessions in rats that received injections of saline or flupenthixol. (C) Anticipatory head entries into the food port in rats that received injections of saline or flupenthixol. (D) Preference ratio in rats receiving saline (left) or flupenthixol (right) injections. (E) Reward consumption for each flavor during the preference tests in rats receiving saline (left) or flupenthixol (right) injections. (F,G) Linear regression relating the change in food consumption as a function of the change in the preference ratio. * *p* < 0.05, ** *p* < 0.01, *** *p* < 0.001.

The OFC encodes value-based parameters and participates in reward-learning and decision-making^19–25^, which suggests the OFC could mediate the change in reward preference. We performed excitotoxic lesions of the OFC or sham surgeries prior to the initial preference test (**Fig. 4A-B**). There was no difference in anticipatory head entries into the food port between sham and OFC-lesioned rats across training sessions (three-way mixed-effects analysis; session effect: *F*_(4, 72)_ = 42.42, *p* = 0.0001; treatment effect: *F*_(1, 18)_ = 1.33, *p* = 0.2648; delay effect: *F*_(1, 18)_ = 0.43, *p* = 0.5213; interaction effect: *F*_(4, 72)_ = 0.25, *p* = 0.9083; **Fig. 4C**). The latency to retrieve the food pellet also did not differ between sham and OFC-lesioned rats across training sessions (three-way mixed-effect analysis; session effect: *F*_(4, 72)_ = 42.42, *p* < 0.0001; treatment effect: *F*_(1, 18)_ = 0.099, *p* = 0.7556; delay effect: *F*_(1, 18)_ = 0.05, *p* = 0.8211; interaction effect: *F*_(4, 72)_ = 0.90, *p* = 0.4697; n = 10 sham rats, n = 10 lesion rats; **Fig. 4D**). Lesioning the OFC did not prevent the change in preference following training (two-way mixed-effects analysis; training effect: *F*_(1, 18)_ = 28.64, *p* < 0.0001; treatment effect: *F*_(1, 18)_ = 0.35, *p* = 0.5599; interaction effect: *F*_(1, 18)_ = 0.0001, *p* = 0.99; **Fig. 4E**). Training resulted in an enhanced preference for the Initial Less Preferred reward in sham surgery and OFC-lesioned rats (post-hoc Sidak’s test; sham: *t*_(18)_ = 3.78, *p* = 0.0028; lesion: *t*_(18)_ = 3.79, *p* = 0.0027; **Fig. 4E**). Both groups exhibited a selective increase in the consumption of the Initial Less Preferred reward (post-hoc Sidak’s test; sham: *t*_(9)_ = 3.89, *p* = 0.0074; lesion: *t*_(9)_ = 3.26, *p* = 0.0195; **Fig. 4F**) with no change in the consumption of the Initial Preferred reward (post-hoc Sidak’s test; sham: *t*_(9)_ = 0.47*, p* = 0.8773; lesion: *t*_(9)_ = 0.73, *p* = 0.7312; **Fig. 4F**). The change in reward preference was positively related to the consumption of the Initial Less Preferred reward (sham: r^2^ = 0.95, *p* < 0.001, lesion: r^2^ = 0.85, *p* < 0.001, **Fig. 4G**) and inversely related to the consumption of the Initial Preferred reward in sham surgery and OFC-lesioned rats (sham: r^2^ = 0.68, *p* < 0.01, lesion: r^2^ = 0.48, *p* = 0.03, **Fig. 4H**). Therefore, the OFC is not involved with the change in reward preference following different delay training sessions.

**Figure 4.**
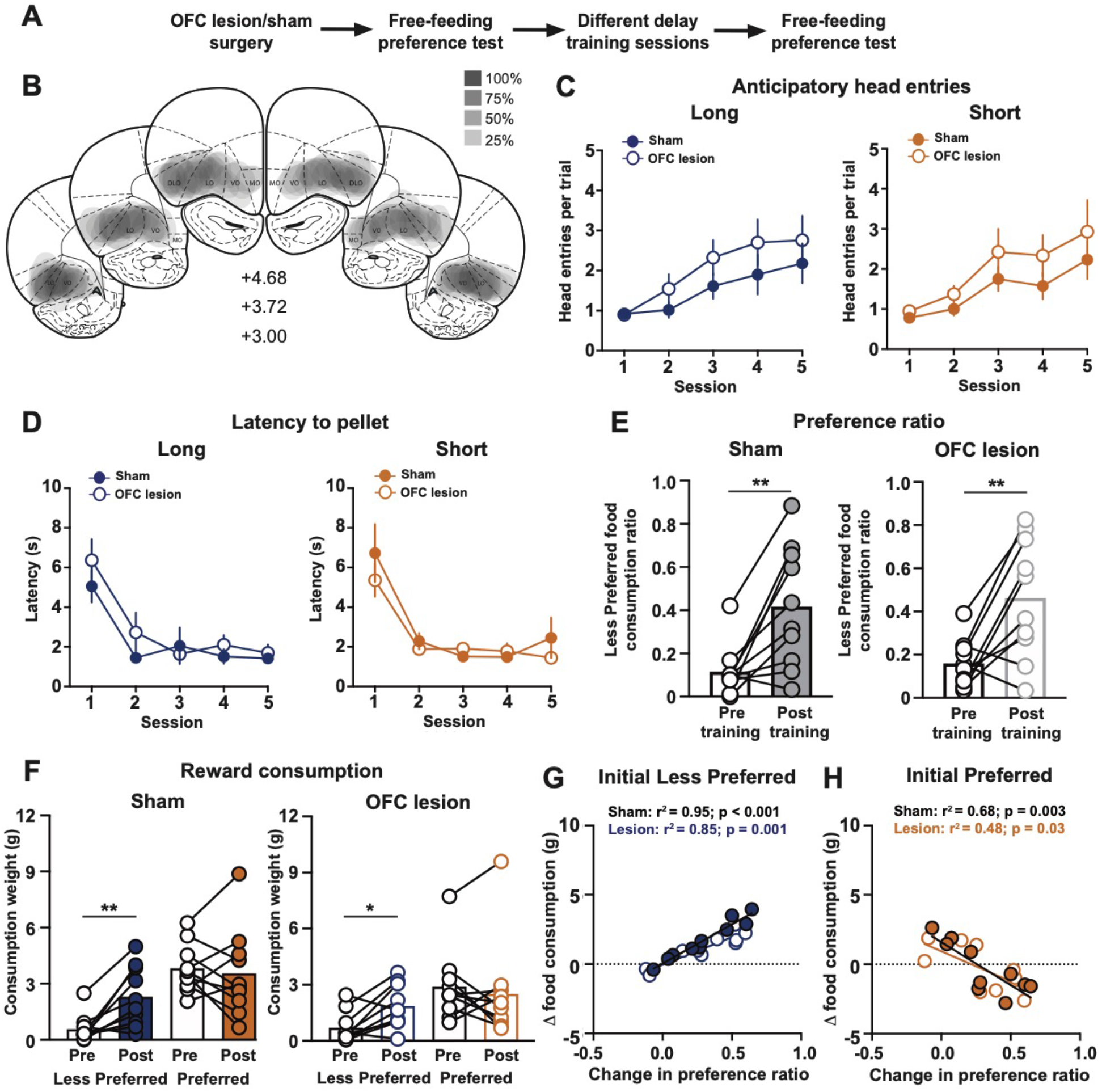
The orbitofrontal cortex (OFC) is not required for the change in reward preference. (A) Training schematic. (B) The extent of OFC lesions across three coronal planes with the anterior distance from bregma (millimeters) indicated. (C) Anticipatory head entries into the food port in sham or OFC-lesioned rats. (D) Latency to make a head entry into the food port after a reward is delivered during training sessions in sham or OFC-lesioned rats. (E) Preference ratio in sham (left) or OFC-lesioned (right) rats. (F) Reward consumption for each flavor during the preference tests in sham (left) or OFC-lesioned (right) rats. (G,H) Linear regression relating the change in food consumption as a function of the change in the preference ratio. * *p* < 0.05, ** *p* < 0.01.

Increasing evidence highlights the BLA is a critical nucleus that contributes to learning, reward valuation, and reward seeking^26–30^. As such, the BLA could potentially mediate the changein reward preference following the different delay training regimen. Rats underwent a surgery to lesion the BLA or a sham procedure prior to the initial preference test (**Fig. 5A-B**). The anticipatory head entries into the food port did not differ between sham and BLA-lesioned rats across training sessions (three-way mixed-effects analysis; session effect: *F*_(2.944, 41.22)_ = 10.45, *p* < 0.0001; treatment effect: *F*_(1, 14)_ = 0.009, *p* = 0.9227; delay effect: *F*_(1, 14)_ = 0.004, *p* = 0.9534; interaction effect: *F*_(4, 56)_ = 0.50, *p* = 0.7384; **Fig. 5C**). There was also no difference in the latency to retrieve a food pellet between sham surgery and BLA-lesioned rats across training sessions (three-way mixed-effect analysis; session effect: *F*_(4, 56)_ = 71.96, *p* < 0.0001; treatment effect: *F*_(1, 14)_ = 0.19, *p* = 0.6702; delay effect: *F*_(1, 14)_ = 0.26, *p* = 0.6190; interaction effect: *F*_(4, 56)_ = 0.05, *p* = 0.9948; n = 8 sham rats, n = 8 lesion rats; **Fig. 5D**). However, lesioning the BLA prevented the change in preference following training (two-way mixed-effects analysis; training effect: *F*_(1, 14)_ = 30.69, *p* < 0.0001; treatment effect: *F*_(1, 14)_ = 3.55, *p* = 0.0805; interaction effect: *F*_(1, 14)_ = 15.17, *p* = 0.0016; **Fig. 5E**). Sham surgery rats exhibited an enhanced preference for the Initial Less Preferred reward (post-hoc Sidak’s test; sham: *t*_(14)_ = 6.67, *p* < 0.0001; **Fig. 5E**) which was due to a selective increase in the consumption of the Initial Less Preferred reward (post-hoc Sidak’s test; Initial Less Preferred flavor: *t*_(7)_ = 5.99, *p* = 0.0011; Initial Preferred flavor: *t*_(7)_ = 2.2, *p* = 0.1233; **Fig. 5F**). In contrast, BLA lesions prevented the change in reward preference (post-hoc Sidak’s test; lesion: *t*_(14)_ = 1.16, *p* = 0.4588; **Fig. 5E**), as rats selectively increased the consumption of the Initial Preferred reward (post-hoc Sidak’s test; Initial Less Preferred flavor: *t*_(7)_ = 2.4, *p* = 0.0931; Initial Preferred flavor: *t*_(7)_ = 4.13, *p* = 0.0088; **Fig. 5F**). The change in reward preference in sham surgery rats was inversely related to the change in consumption of the Initial Preferred reward (Initial Less Preferred: r^2^ = 0.02, *p* = 0.14; Initial Preferred: r^2^ = 0.60, *p* = 0.02; **Fig. 5G-H**). A similar trend was observed in BLA-lesioned rats (Initial Less Preferred: r^2^ = 0.38, *p* = 0.1; Initial Preferred: r^2^ = 0.52, *p* = 0.04; **Fig. 5G-H**). These data collectively illustrate that the BLA is required for the change in reward preference elicited by sunk temporal costs.

**Figure 5.**
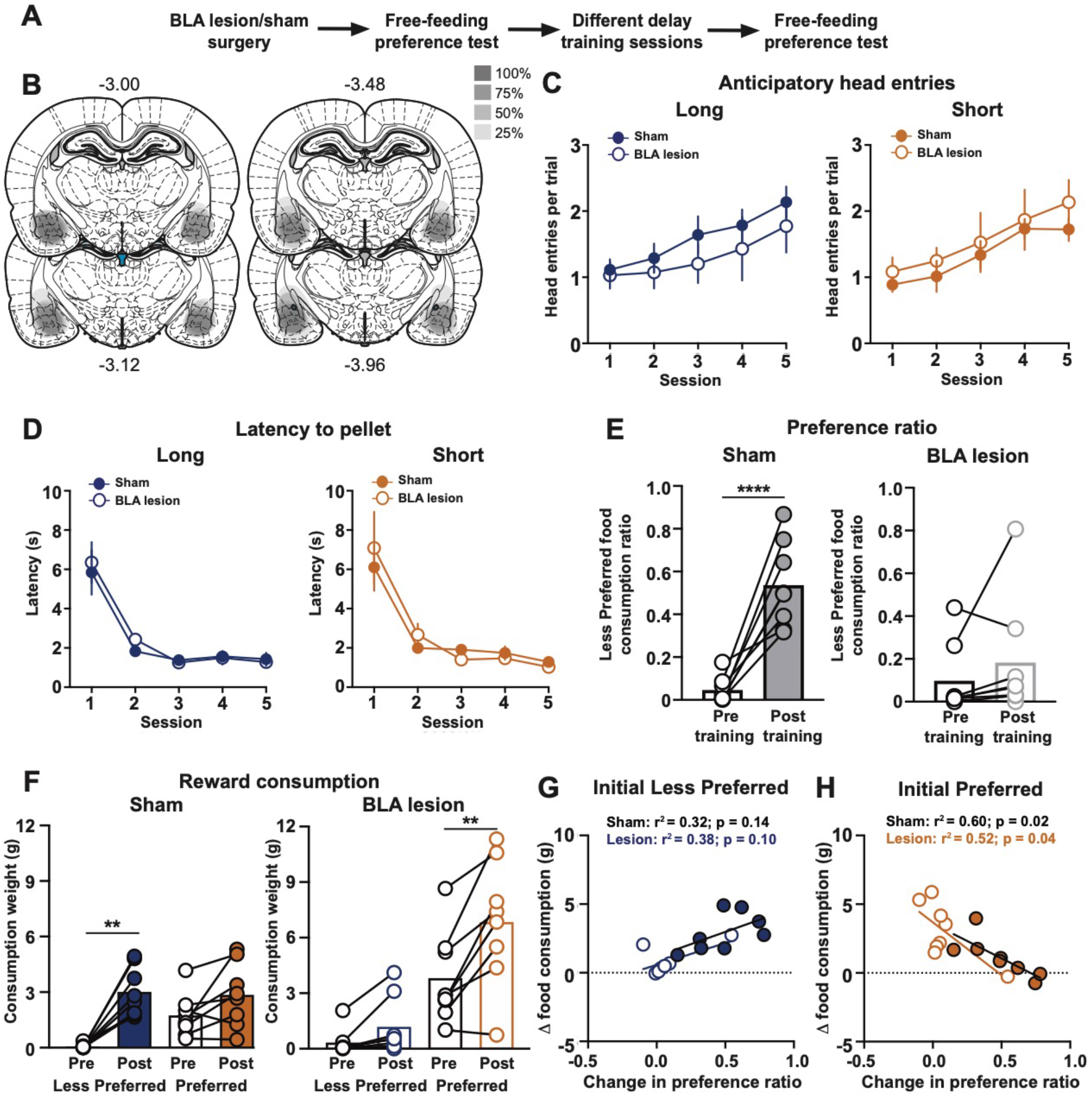
The basolateral amygdala (BLA) is required for the change in reward preference. (A) Training schematic. (B) The extent of BLA lesions across four coronal planes with the anterior distance from bregma (millimeters) indicated. (C) Anticipatory head entries into the food port in sham or BLA-lesioned rats. (D) Latency to make a head entry into the food port after a reward is delivered during training sessions in sham or BLA-lesioned rats. (E) Preference ratio in sham (left) or BLA-lesioned (right) rats. (F) Reward consumption for each flavor during the preference tests in sham (left) or BLA-lesioned (right) rats. (G,H) Linear regression relating the change in food consumption as a function of the change in the preference ratio. * *p* < 0.05, ** *p* < 0.01, *** *p* < 0.001.

The RSC is another brain region that could participate in sustained changes in reward preference given its role in learning, and that RSC neurons respond to rewards to encode value-related signals^32–35^. To address this possibility, rats underwent an RSC lesion or a sham surgery prior to the initial preference test (**Fig. 6A-B**). We found a significant interaction between treatment and delay on the anticipatory head entries between sham and RSC-lesioned rats (three-way mixed-effects analysis; session effect: *F*_(3.086, 43.21)_ = 14.14, *p* < 0.0001; treatment effect: *F*_(1, 53)_ = 0.10, *p* = 0.7504; delay effect: *F*_(1, 14)_ = 0.06, *p* = 0.8134; treatment x delay effect: *F*_(1, 53)_ = 5.04, *p* = 0.029; interaction effect: *F*_(4, 53)_ = 2.44, *p* = 0.058; **Fig. 6C**). There was no difference in the post-reward latency to retrieve a food pellet between sham surgery and RSC-lesioned rats across training sessions (three-way mixed-effect analysis; session effect: *F*_(4, 56)_ = 43.99, *p* < 0.0001; treatment effect: *F*_(1, 53)_ = 0.18, *p* = 0.6756; delay effect: *F*_(1, 14)_ = 0.59, *p* = 0.4543; interaction effect: *F*_(4, 53)_ = 1.26, *p* = 0.2952; n = 8 sham rats, n = 8 lesion rats; **Fig. 6D**). Lesioning the RSC prevented the change in preference following training (two-way mixed-effects analysis; training effect: *F*_(1, 14)_ = 10.64, *p* = 0.0057; treatment effect: *F*_(1, 14)_ = 12.69, *p* = 0.0031; interaction effect: *F*_(1, 14)_ = 17.36, *p* = 0.001; **Fig. 6E**). Sham rats exhibited an enhanced preference for the Initial Less Preferred reward (post-hoc Sidak’s test; sham: *t*_(14)_ = 5.25, *p* = 0.0002; **Fig. 6E**), which was accompanied by an increased consumption of both rewards following training (post-hoc Sidak’s test; Initial Less Preferred flavor: *t*_(7)_ = 4.31, *p* = 0.0071; Initial Preferred flavor: *t*_(7)_ = 2.95, *p* = 0.0424; **Fig. 6F**). In contrast, there was no change in preference in the RSC lesion group (post-hoc Sidak’s test; lesion: *t*_(14)_ = 0.64, *p* = 0.7814; **Fig. 6E**), as rats selectively increased the consumption of the Initial Preferred reward (post-hoc Sidak’s test; Initial Less Preferred flavor: *t*_(7)_ = 0.28, *p* = 0.9548; Initial Preferred flavor: *t*_(7)_ = 10.63, *p* < 0.0001; **Fig. 6F**). The relative change in reward preference and consumption of the Initial Less Preferred reward was positively correlated across RSC lesion and sham groups (Initial Less Preferred sham: r^2^ = 0.52, *p* = 0.04, lesion: r^2^ = 0.87, *p* < 0.001; Initial Preferred sham: r^2^ = 0.02, *p* = 0.74; lesion: r^2^ = 0.36, *p* = 0.11; **Fig. 6G-H**). Our results highlight that both the BLA and RSC are necessary for increasing the preference toward an initial less desirable reward option.

**Figure 6.**
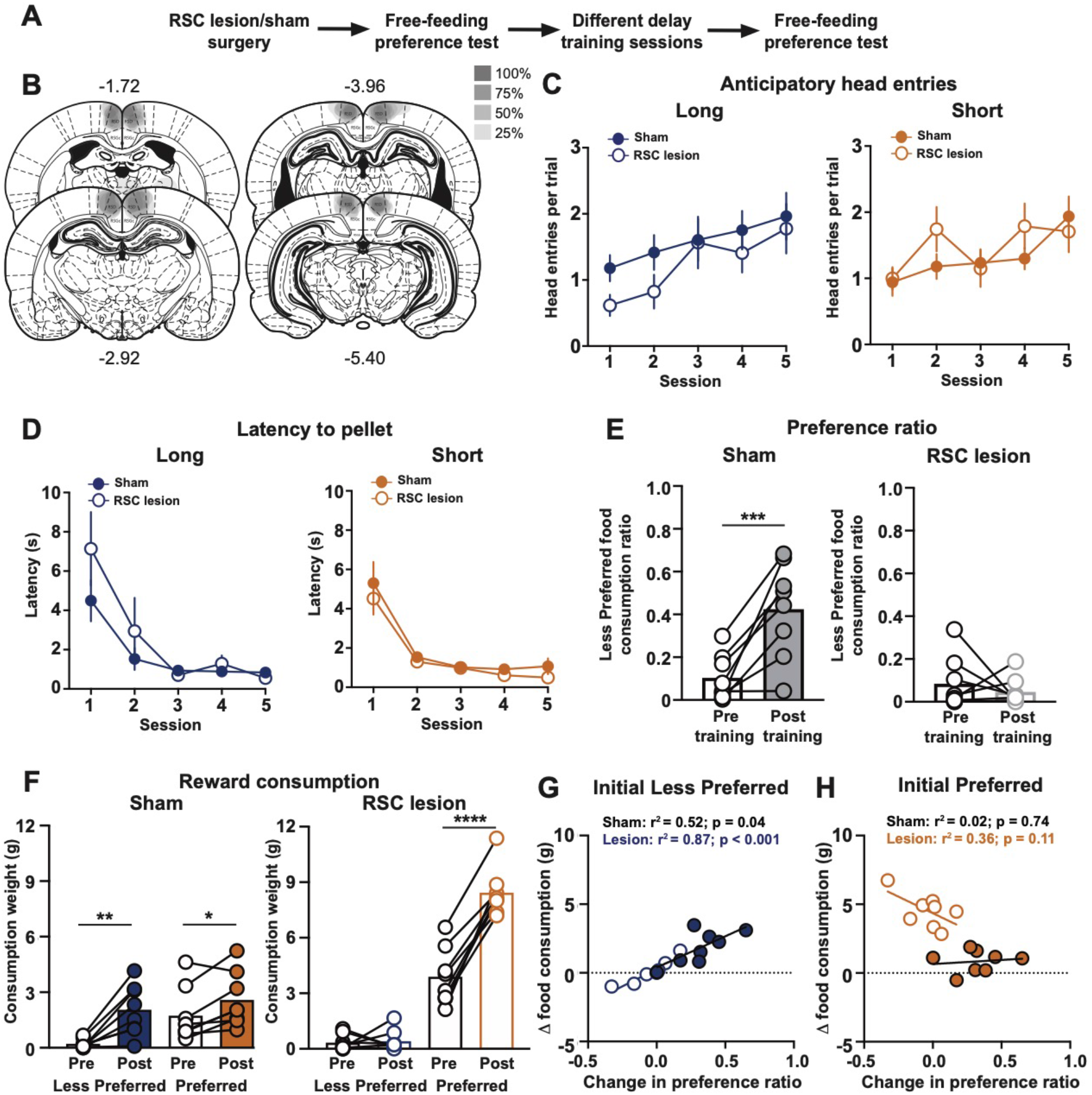
The retrosplenial cortex (RSC) is required for the change in reward preference. (A) Training schematic. (B) The extent of RSC lesions across four coronal planes with the anterior distance from bregma (millimeters) indicated. (C) Anticipatory head entries into the food port in sham or RSC-lesioned rats. (D) Latency to make a head entry into the food port after a reward is delivered during training sessions in sham or RSC-lesioned rats. (E) Preference ratio in sham (left) or RSC-lesioned (right) rats. (F) Reward consumption for each flavor during the preference tests in sham (left) or RSC-lesioned (right) rats. (G,H) Linear regression relating the change in food consumption as a function of the change in the preference ratio. * *p* < 0.05, ** *p* < 0.01, *** *p* < 0.001, \*\*\*\**p* < 0.0001.

When examining all subjects trained on the different delay regimen, the change in reward preference was positively related to the consumption of the Initial Less Preferred reward (r^2^ = 0.76, *p* < 0.001) and inversely related to the consumption of the Initial Preferred reward (r^2^ = 0.47, *p* < 0.001; n = 88 rats). This suggests that rats exhibiting a robust bias toward the Initial Less Preferred reward could potentially decrease the consumption of the Initial Preferred reward following training. We therefore analyzed the relative food consumption across subjects based upon if there was a mild bias toward the Initial Preferred reward (**Fig. 7A**, orange), a mild bias toward the Initial Less Preferred reward (**Fig. 7A**, light blue), or a robust bias toward the Initial Less Preferred reward (**Fig. 7A**, dark blue). The relative change in food consumption for each reward differed according to the change in reward preference (two-way mixed-effects analysis; flavor effect: *F*_(1, 85)_ = 3.89, *p* = 0.0517; change in preference effect: *F*_(2, 85)_ = 5.78, *p* = 0.0044; interaction effect: *F*_(2, 85)_ = 116.2, *p* < 0.0001; **Fig. 7B-D**). Rats exhibiting a mild bias toward the original reward preference selectively increased the consumption of the Initial Preferred reward (one-sample t-test relative to 0: Initial Less Preferred: *t*_(15)_ = 0.69, *p* = 0.5007; Initial Preferred: *t*_(15)_ = 8.39, *p* < 0.0001; **Fig. 7B**). Rats with a mild bias toward the Initial Less Preferred reward increased the consumption of both rewards (one-sample t-test relative to 0: Initial Less Preferred: *t*_(44)_ = 88.88*, p* < 0.0001; Initial Preferred: *t*_(44)_ = 5.47*, p* < 0.0001; **Fig. 7C**). Interestingly, rats with a robust bias toward the Initial Less Preferred reward increased the consumption of the Initial Less Preferred and *decreased* the consumption of the Initial Preferred reward (one-sample t-test relative to 0: Initial Less Preferred: *t*_(26)_ = 16.99*, p* < 0.0001; Initial Preferred: *t*_(26)_ = 2.23*, p* = 0.0348; **Fig. 7D**). Together, these results highlight that sunk temporal costs toward one reward can positively impact the value of that reward as well as negatively impact the value of the alternative option.

**Figure 7.**
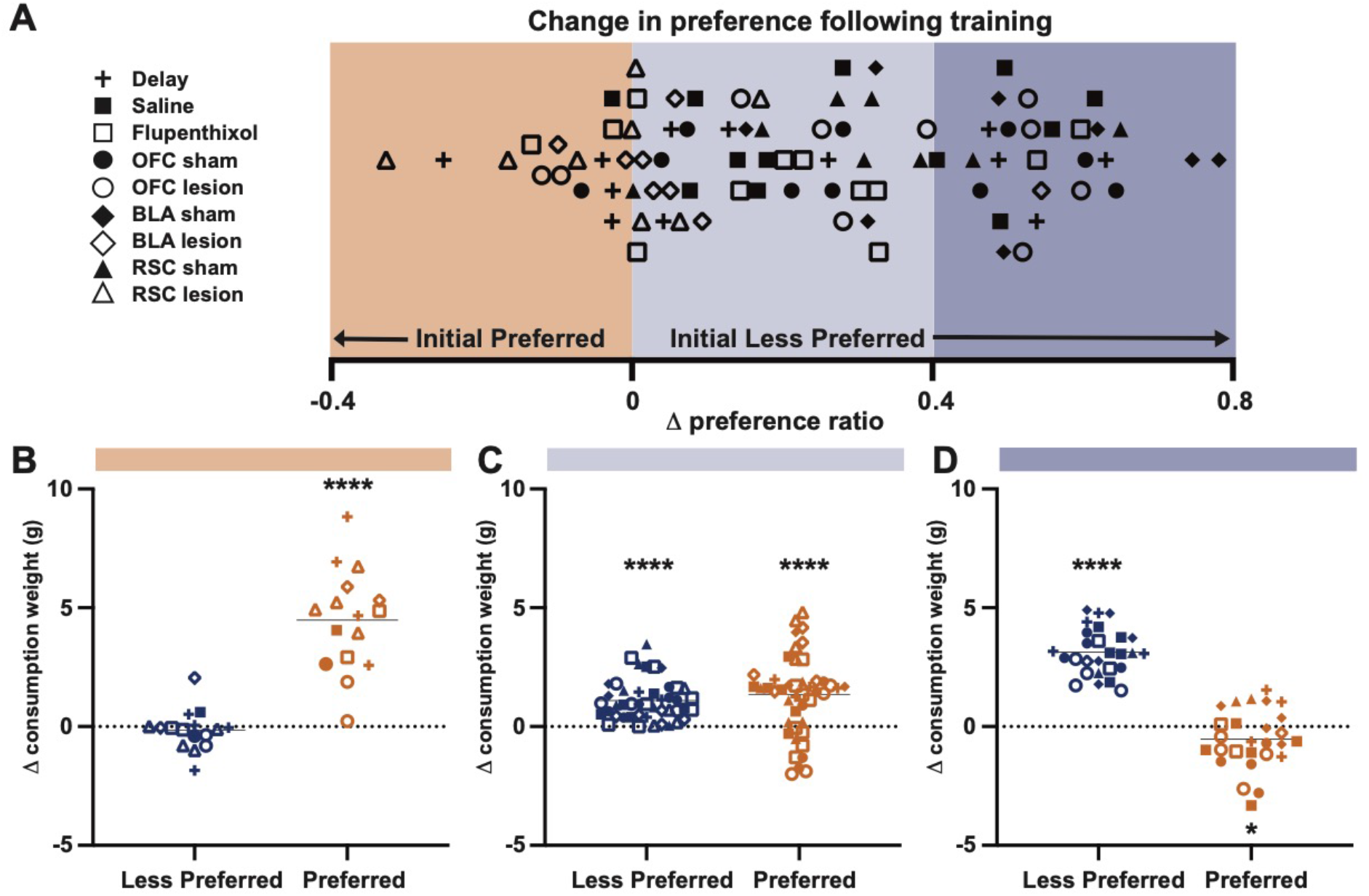
Relating the change in preference to the change in the food consumption. (A) Change in the preference ratio across all rats that underwent the different delay training sessions. Color overlays denote a mild bias toward the Initial Preferred reward (orange; change in the preference ratio < 0), a mild bias toward the Initial Less Preferred reward (light blue; change in the preference ratio between 0 and 0.4), and a robust bias toward the Initial Less Preferred reward (dark blue; change in the preference ratio > 0.4). (B) Change in the food consumption in rats that displayed a mild bias toward the Initial Preferred reward. (C) Change in the food consumption in rats that displayed a mild bias toward the Initial Less Preferred reward. (D) Change in the food consumption in rats that displayed a robust bias toward the Initial Less Preferred reward. * *p* < 0.05, **** *p* < 0.0001.

## Discussion

Sunk costs are irrecoverable past costs that can influence future decisions. Human studies illustrate behavioral consequences from a variety of sunk costs that are imposed on the subject, including embarrassment, political, personal, and financial, though these costs are challenging to model in animals^10,12,39–41^. Research in pigeons and humans demonstrate that subjects will display increased preference for an outcome that follows greater effort^5–7,9^. However, increasing effort (e.g. more responses) often requires more time, thereby confounding effort costs with temporal costs^42^. As such, exclusively manipulating temporal costs provides an attractive strategy to examine the impact of sunk costs across species^11,43^. Recent studies have examined the behavioral impact of sunk temporal costs using a choice-based foraging task^11,43,44^. In this foraging task, choosing one option comes at the expense of a potential alternative outcome, so one cannot determine if sunk costs increase the value of the chosen outcome and/or decrease the value of the alternative outcome. To determine how sunk temporal costs impact reward value, we quantified the change in reward consumption and reward preference using a free-feeding assay. We found that sunk temporal costs resulted in increased consumption of the Initial Less Preferred reward after training. Interestingly, we observed that a robust change in preference toward the Initial Less Preferred reward were a product of changes in consumption of *both* rewards: increased consumption of the Initial Less Preferred reward as well as decreased consumption of the Initial Preferred reward.

Together, these findings indicate that sunk temporal costs toward one reward can positively impact the value of that reward as well as negatively impact the value of the alternative option.

While our training paradigm was designed to examine the impact of sunk temporal costs, it is important to address potential alternative factors that could contribute to our behavioral findings. In control experiments we found that rats maintained their initial reward preference when the delay to the reward delivery was held constant for both flavors. These results illustrate that rats are appropriately discriminating between the reward options within the task design. The change in preference is not due to satiety since rats underwent only a single training session per day and the same number of food pellets were delivered in each training session. With this experimental design, the reward density is lower in the Long Delay sessions relative to the Short Delay sessions. As such, the pellet delivery in the Long Delay sessions could be perceived as an unexpected reward. However, this interpretation is not supported by our data as there was no difference in anticipatory responding prior to the reward delivery between Long and Short Delay sessions.

Prior research has identified a number of factors that can alter reward preference. For example, allowing free access to one reward option prior to a preference test will devalue that reward and elicit a transient change in preference^45–47^. Long term changes in preference can be induced by pairing a reward with an aversive outcome^48^. Here, we utilized a long temporal delay before the delivery of an initial less desirable option to subsequently increase preference for that reward. Critically, the post-training preference test is performed in a distinct context and on the day following the last training session. Taken together, these results demonstrate that the temporal costs imposed during training sessions can induce a sustained change in preference without the need for altering satiety or pairing the outcome with an aversive stimulus.

Dopamine signals reflect changes in preference induced by state-specific satiety^46,47^. Additionally, dopamine receptor antagonism impairs preference changes arising from conditioned taste aversion^49^. In our task, we found that antagonizing dopamine receptors during training sessions increased the latency to retrieve the reward and decreased anticipatory head entries into the food port, consistent with dopamine’s role in regulating both locomotor activity and the acquisition of anticipatory responding^37,50,51^. Despite the motoric effects of dopamine receptor antagonism during training sessions, rats still displayed enhanced preference for the initial less desirable reward. Therefore, behavioral performance during training sessions is not linked to a subsequent change in reward preference.

We additionally found that OFC lesions failed to prevent changes in reward preference. These results are consistent with previous reports demonstrating that lesions to the OFC did not alter outcome preference or behavioral flexibility^52,53^. However, prior research has found that OFC lesions impair the ability to update choices following selective satiation^23^. In addition, OFC lesions do not prevent changes in preference induced by taste aversion, but can disrupt devaluation of the associated cue^54^. While the dopamine system and the OFC are active participants in other forms of preference changes, our findings suggest these two systems do not regulate changes in preference elicited by sunk temporal costs.

Our results demonstrate that lesioning the BLA prior to training sessions prevented sunk temporal costs from influencing changes in preference. Previous studies have implicated BLA neurons in reward learning and memory formation^26,27^. However, the BLA also has a role in reward preference as studies demonstrate the BLA maintains representations of appetitive and economic value^30,55,56^. BLA neurons additionally encode the value of a chosen reward and inactivating the BLA decreased choice for a more preferred option^57,58^. Moreover, BLA inactivation impairs choice following reinforcer devaluation^58,59^. Our findings coupled with prior work collectively highlight a critical role for the BLA in updating reward value.

The RSC has been well-studied for its contributions to memory formation, spatial navigation, and timing^32,60–63^. However, increasing evidence illustrates that the RSC also contains reward-responsive neurons that encode reward value^33–35^. Furthermore, inactivation of the RSC impaired the ability to adapt behavior based on the reward history^34^. Consistent with the RSC’s role in reward-based behavior, RSC lesions prior to training sessions prevented the change in preference induced by sunk temporal costs. RSC lesions also differentially altered the anticipatory responding between training sessions. However, as discussed above, the behavioral responses during training sessions are not predictive of changes in reward preference.

Lesions of the BLA or the RSC prevented the change in reward preference. These results indicate the presence of a circuit involving the BLA and RSC to update reward value. In support, the BLA sends direct projections to the RSC^64^. We note that the lesion experiments demonstrate the necessity of the BLA and RSC in changing reward preference. However, future studies are needed to determine if the BLA and RSC are responsible for acquiring and/or expressing changes in reward preference. Taken together, our data highlights a previously unappreciated role for the BLA and RSC in mediating changes in preference following sunk temporal costs.

## Acknowledgements

This work was supported by National Institutes of Health grants DA033386 (MJW) and DA042362 (MJW) and the Mind Science Foundation Research Award (MJL). The authors declare no conflict of interest.

## Author Contributions

MJL, APM, JMG, MRL performed the experiments and analyzed the data. MJL and MJW designed the experiments and wrote the manuscript.

