## Supplementary Data for "Enhanced preference for delayed rewards requires the basolateral amygdala and retrosplenial cortex"

| Figure 1 – Different delay training | | | |
|---|---|---|---|
| **Panel B – Anticipatory head entries** | | | |
| Two-way mixed-effects model | Session<br>$F_{(1.894, 20.83)} = 6.52$, $\boldsymbol{p = 0.007}$ | Delay<br>$F_{(1, 11)} = 0.05$, $p = 0.8337$ | Two-way interaction<br>$F_{(2.661, 29.27)} = 0.92$, $p = 0.4334$ |
| **Panel C – Latency to pellet** | | | |
| Two-way mixed-effects model | Session<br>$F_{(1.316, 14.48)} = 54.17$, $\boldsymbol{p < 0.0001}$ | Delay<br>$F_{(1, 11)} = 0.16$, $p = 0.7002$ | Two-way interaction<br>$F_{(1.30, 12.43)} = 0.23$, $p = 0.6672$ |
| **Panel D – Preference ratio** | | | |
| Paired t-test | | $t_{(11)} = 2.33$, $\boldsymbol{p = 0.0402}$ | |
| **Panel E – Reward consumption** | | | |
| Two-way mixed-effects model | Training<br>$F_{(1, 11)} = 15.56$, $\boldsymbol{p = 0.0023}$ | Flavor<br>$F_{(1, 11)} = 12.27$, $\boldsymbol{p = 0.0049}$ | Two-way interaction<br>$F_{(1, 11)} = 0.99$, $p = 0.3417$ |
| Post-hoc Sidak's test | Initial Less Preferred<br>$t_{(11)} = 2.65$, $\boldsymbol{p = 0.0451}$ | Initial Preferred<br>$t_{(11)} = 2.98$, $\boldsymbol{p = 0.0251}$ | |
| **Panel F – Initial Less Preferred correlation** | $r^2 = 0.96$, $\boldsymbol{p < 0.0001}$ | **Panel G – Initial Preferred correlation** | $r^2 = 0.38$, $\boldsymbol{p = 0.03}$ |

| Figure 2 – Same delay training | | | |
|---|---|---|---|
| **Panel B – Anticipatory head entries** | | | |
| Two-way mixed-effects model | Session<br>$F_{(2.831, 16.99)} = 37.89$, $\boldsymbol{p < 0.0001}$ | Delay<br>$F_{(1, 6)} = 1.20$, $p = 0.3146$ | Two-way interaction<br>$F_{(2.297, 13.78)} = 1.58$, $p = 0.2414$ |
| **Panel C – Latency to pellet** | | | |
| Two-way mixed-effects model | Session<br>$F_{(1.341, 8.049)} = 2.97$, $p = 0.1178$ | Delay<br>$F_{(1, 6)} = 14.93$, $p = 0.2675$ | Two-way interaction<br>$F_{(2.094, 12.04)} = 0.62$, $p = 0.5636$ |
| **Panel D – Preference ratio** | | | |
| Paired t-test | | $t_{(6)} = 0.91$, $p = 0.40$ | |
| **Panel E – Reward consumption** | | | |
| Two-way mixed-effects model | Training<br>$F_{(1, 6)} = 15.15$, $\boldsymbol{p = 0.008}$ | Flavor<br>$F_{(1, 6)} = 23.47$, $\boldsymbol{p = 0.0029}$ | Two-way interaction<br>$F_{(1, 6)} = 3.99$, $p = 0.0926$ |
| Post-hoc Sidak's test | Initial Less Preferred<br>$t_{(6)} = 1.56$, $p = 0.3104$ | Initial Preferred<br>$t_{(6)} = 3.71$, $\boldsymbol{p = 0.0199}$ | |
| **Panel F – Initial Less Preferred correlation** | $r^2 = 0.16$, $p = 0.38$ | **Panel G – Initial Preferred correlation** | $r^2 = 0.00$, $p = 0.97$ |

| Figure 3 – Saline vs Flupenthixol | | | |
|---|---|---|---|
| **Panel B – Anticipatory head entries** | | | |
| Three-way mixed-effects model | Session<br>$F_{(3.020, 66.43)} = 22.26$, $\boldsymbol{p < 0.0001}$ | Treatment<br>$F_{(1, 87)} = 11.22$, $\boldsymbol{p = 0.0012}$ | Delay<br>$F_{(1, 22)} = 0.05$, $p = 0.8188$ |
| Session x Treatment<br>$F_{(4, 87)} = 11.22$, $\boldsymbol{p < 0.0001}$ | Session x Delay<br>$F_{(3.144, 66.37)} = 0.14$, $p = 0.9405$ | Treatment x Delay<br>$F_{(1, 87)} = 1.88$, $p = 0.1743$ | Three-way interaction<br>$F_{(4, 87)} = 0.32$, $p = 0.8644$ |
| **Panel C – Latency to pellet** | | | |
| Three-way mixed-effects model | Session<br>$F_{(4, 88)} = 43.04$, $\boldsymbol{p < 0.0001}$ | Treatment<br>$F_{(1, 22)} = 6.18$, $\boldsymbol{p = 0.016}$ | Delay<br>$F_{(1, 22)} = 0.02$, $p = 0.8840$ |
| Session x Treatment<br>$F_{(4, 88)} = 0.39$, $p = 0.8167$ | Session x Delay<br>$F_{(4, 88)} = 1.14$, $p = 0.3454$ | Treatment x Delay<br>$F_{(1, 22)} = 0.0007$, $p = 0.9781$ | Three-way interaction<br>$F_{(4, 88)} = 1.53$, $p = 0.2010$ |
| **Panel D – Preference ratio** | | | |
| Two-way mixed-effects model | Training<br>$F_{(1, 22)} = 30.86$, $\boldsymbol{p < 0.0001}$ | Treatment<br>$F_{(1, 22)} = 0.11$, $p = 0.7384$ | Two-way interaction<br>$F_{(1, 22)} = 0.77$, $p = 0.3901$ |
| Post-hoc Sidak's test | Saline<br>$t_{(22)} = 4.55$, $\boldsymbol{p = 0.0003}$ | Flupenthixol<br>$t_{(22)} = 3.31$, $\boldsymbol{p = 0.0064}$ | |
| **Panel E – Reward consumption** | | | |
| Saline<br>Two-way mixed-effects model | Training<br>$F_{(1, 11)} = 12.38$, $\boldsymbol{p = 0.0048}$ | Flavor<br>$F_{(1, 11)} = 11.75$, $\boldsymbol{p = 0.0056}$ | Two-way interaction<br>$F_{(1, 11)} = 4.04$, $p = 0.0697$ |
| Post-hoc Sidak's test | Initial Less Preferred<br>$t_{(11)} = 4.92$, $\boldsymbol{p = 0.0009}$ | Initial Preferred<br>$t_{(11)} = 0.91$, $p = 0.6169$ | |

| | | | |
|---|---|---|---|
| Flupenthixol<br>Two-way mixed-effects model | Training<br>$F_{(1, 11)} = 12$, $\mathbf{p = 0.0053}$ | Flavor<br>$F_{(1, 11)} = 14.05$, $\mathbf{p = 0.0032}$ | Two-way interaction<br>$F_{(1, 11)} = 0.30$, $p = 0.5930$ |
| Post-hoc Sidak's test | Initial Less Preferred<br>$t_{(11)} = 3.54$, $\mathbf{p = 0.0092}$ | Initial Preferred<br>$t_{(11)} = 1.77$, $p = 0.1984$ | |
| **Panel F – Initial Less Preferred correlation** | Saline<br>$r^2 = 0.72$, $\mathbf{p < 0.001}$ | Flupenthixol<br>$r^2 = 0.74$, $\mathbf{p < 0.001}$ | |
| **Panel G – Initial Preferred correlation** | Saline<br>$r^2 = 0.54$, $\mathbf{p < 0.01}$ | Flupenthixol<br>$r^2 = 0.46$, $\mathbf{p = 0.02}$ | |

| **Figure 4 – OFC lesion vs sham** | | | |
|---|---|---|---|
| **Panel C – Anticipatory head entries** | | | |
| Three-way mixed-effects model | Session<br>$F_{(4, 72)} = 42.42$, $\mathbf{p = 0.0001}$ | Treatment<br>$F_{(1, 18)} = 1.33$, $p = 0.2648$ | Delay<br>$F_{(1, 18)} = 0.43$, $p = 0.5213$ |
| Session x Treatment<br>$F_{(4, 72)} = 0.58$, $p = 0.6799$ | Session x Delay<br>$F_{(2.363, 42.54)} = 1.86$, $p = 0.1614$ | Treatment x Delay<br>$F_{(1, 18)} = 0.009$, $p = 0.9238$ | Three-way interaction<br>$F_{(4, 72)} = 0.25$, $p = 0.9083$ |
| **Panel D – Latency to pellet** | | | |
| Three-way mixed-effects model | Session<br>$F_{(4, 72)} = 42.42$, $\mathbf{p < 0.0001}$ | Treatment<br>$F_{(1, 18)} = 0.099$, $p = 0.7556$ | Delay<br>$F_{(1, 18)} = 0.05$, $p = 0.8211$ |
| Session x Treatment<br>$F_{(4, 72)} = 0.39$, $p = 0.8188$ | Session x Delay<br>$F_{(4, 72)} = 0.14$, $p = 0.9665$ | Treatment x Delay<br>$F_{(1, 18)} = 1.87$, $p = 0.1879$ | Three-way interaction<br>$F_{(4, 72)} = 0.90$, $p = 0.4697$ |
| **Panel E – Preference ratio** | | | |
| Two-way mixed-effects model | Training<br>$F_{(1, 18)} = 28.64$, $\mathbf{p < 0.0001}$ | Treatment<br>$F_{(1, 18)} = 0.35$, $p = 0.5599$ | Two-way interaction<br>$F_{(1, 18)} = 0.0001$, $p = 0.99$ |
| Post-hoc Sidak's test | Sham<br>$t_{(18)} = 3.78$, $\mathbf{p = 0.0028}$ | OFC lesion<br>$t_{(18)} = 3.79$, $\mathbf{p = 0.0027}$ | |
| **Panel F – Reward consumption** | | | |
| Sham<br>Two-way mixed-effects model | Training<br>$F_{(1, 9)} = 3.82$, $\mathbf{p = 0.0825}$ | Flavor<br>$F_{(1, 9)} = 14.1$, $\mathbf{p = 0.0045}$ | Two-way interaction<br>$F_{(1, 9)} = 7.17$, $\mathbf{p = 0.0253}$ |
| Post-hoc Sidak's test | Initial Less Preferred<br>$t_{(9)} = 3.89$, $\mathbf{p = 0.0074}$ | Initial Preferred<br>$t_{(9)} = 0.47$, $p = 0.8773$ | |
| OFC lesion<br>Two-way mixed-effects model | Training<br>$F_{(1, 9)} = 1.56$, $p = 0.2431$ | Flavor<br>$F_{(1, 9)} = 4.73$, $\mathbf{p = 0.0577}$ | Two-way interaction<br>$F_{(1, 9)} = 6.04$, $p = 0.0363$ |
| Post-hoc Sidak's test | Initial Less Preferred<br>$t_{(9)} = 3.26$, $\mathbf{p = 0.0195}$ | Initial Preferred<br>$t_{(9)} = 0.73$, $p = 0.7312$ | |
| **Panel G – Initial Less Preferred correlation** | Sham<br>$r^2 = 0.95$, $\mathbf{p < 0.001}$ | OFC lesion<br>$r^2 = 0.85$, $\mathbf{p < 0.001}$ | |
| **Panel H – Initial Preferred correlation** | Sham<br>$r^2 = 0.68$, $\mathbf{p < 0.01}$ | OFC lesion<br>$r^2 = 0.48$, $\mathbf{p = 0.03}$ | |

| **Figure 5 – BLA lesion vs sham** | | | |
|---|---|---|---|
| **Panel C – Anticipatory head entries** | | | |
| Three-way mixed-effects model | Session<br>$F_{(2.944, 41.22)} = 10.45$, $\mathbf{p < 0.0001}$ | Treatment<br>$F_{(1, 14)} = 0.009$, $p = 0.9227$ | Delay<br>$F_{(1, 14)} = 0.004$, $p = 0.9534$ |
| Session x Treatment<br>$F_{(4, 56)} = 0.12$, $p = 0.9732$ | Session x Delay<br>$F_{(2.88, 40.33)} = 0.67$, $p = 0.5686$ | Treatment x Delay<br>$F_{(1, 14)} = 3.80$, $p = 0.0717$ | Three-way interaction<br>$F_{(4, 56)} = 0.50$, $p = 0.7384$ |
| **Panel D – Latency to pellet** | | | |
| Three-way mixed-effects model | Session<br>$F_{(4, 56)} = 71.96$, $\mathbf{p < 0.0001}$ | Treatment<br>$F_{(1, 14)} = 0.19$, $p = 0.6702$ | Delay<br>$F_{(1, 14)} = 0.26$, $p = 0.6190$ |
| Session x Treatment<br>$F_{(4, 56)} = 1.02$, $p = 0.4057$ | Session x Delay<br>$F_{(4, 56)} = 0.14$, $p = 0.9678$ | Treatment x Delay<br>$F_{(1, 14)} = 0.001$, $p = 0.9736$ | Three-way interaction<br>$F_{(4, 56)} = 0.05$, $p = 0.9948$ |
| **Panel E – Preference ratio** | | | |
| Two-way mixed-effects model | Training<br>$F_{(1, 14)} = 30.69$, $\mathbf{p < 0.0001}$ | Treatment<br>$F_{(1, 14)}$ 3.55, $p = 0.0805$ | Two-way interaction<br>$F_{(1, 14)} = 15.17$, $\mathbf{p = 0.0016}$ |
| Post-hoc Sidak's test | Sham<br>$t_{(14)} = 6.67$, $\mathbf{p < 0.0001}$ | BLA lesion<br>$t_{(14)} = 1.16$, $p = 0.4588$ | |
| **Panel F – Reward consumption** | | | |

| | | | |
|---|---|---|---|
| Sham<br>Two-way mixed-effects model | Training<br>$F_{(1,7)}$ = 33.08, ***p* = 0.0007** | Flavor<br>$F_{(1, 7)}$ = 3.53, *p* = 0.1025 | Two-way interaction<br>$F_{(1, 7)}$ = 6.72, ***p* = 0.0358** |
| Post-hoc Sidak's test | Initial Less Preferred<br>$t_{(7)}$ = 5.99, ***p* = 0.0011** | Initial Preferred<br>$t_{(7)}$ = 2.2, *p* = 0.1233 | |
| BLA lesion<br>Two-way mixed-effects model | Training<br>$F_{(1, 7)}$ = 22.63, ***p* = 0.0021** | Flavor<br>$F_{(1, 7)}$ = 18.66, ***p* = 0.0035** | Two-way interaction<br>$F_{(1, 7)}$ = 7.16, ***p* = 0.0317** |
| Post-hoc Sidak's test | Initial Less Preferred<br>$t_{(7)}$ = 2.4, *p* = 0.0931 | Initial Preferred<br>$t_{(7)}$ = 4.13, ***p* = 0.0088** | |
| **Panel G – Initial Less Preferred correlation** | Sham<br>$r^2$ = 0.32, *p* = 0.14 | BLA lesion<br>$r^2$ = 0.38, *p* = 0.1 | |
| **Panel H – Initial Preferred correlation** | Sham<br>$r^2$ = 0.60, ***p* = 0.02** | BLA lesion<br>$r^2$ = 0.52, ***p* = 0.04** | |

| **Figure 6 – RSC lesion vs sham** | | | |
|---|---|---|---|
| **Panel C – Anticipatory head entries** | | | |
| Three-way mixed-effects model | Session<br>$F_{(3.086, 43.21)}$ = 14.14, ***p* < 0.0001** | Treatment<br>$F_{(1, 53)}$ = 0.10, *p* = 0.7504 | Delay<br>$F_{(1, 14)}$ = 0.06, *p* = 0.8134 |
| Session x Treatment<br>$F_{(4, 53)}$ = 0.53, *p* = 0.7152 | Session x Delay<br>$F_{(2.213, 29.32)}$ = 2.29, *p* = 0.1145 | Treatment x Delay<br>$F_{(1, 53)}$ = 5.04, ***p* = 0.029** | Three-way interaction<br>$F_{(4, 53)}$ = 2.44, *p* = 0.058 |
| **Panel D – Latency to pellet** | | | |
| Three-way mixed-effects model | Session<br>$F_{(4, 56)}$ = 43.99, ***p* < 0.0001** | Treatment<br>$F_{(1, 53)}$ = 0.18, *p* = 0.6756 | Delay<br>$F_{(1, 14)}$ = 0.59, *p* = 0.4543 |
| Session x Treatment<br>$F_{(4, 53)}$ = 0.88, *p* = 0.4816 | Session x Delay<br>$F_{(4, 53)}$ = 0.62, *p* = 0.6474 | Treatment x Delay<br>$F_{(1, 53)}$ = 1.64, *p* = 0.2216 | Three-way interaction<br>$F_{(4, 53)}$ = 1.26, *p* = 0.2952 |
| **Panel E – Preference ratio** | | | |
| Two-way mixed-effects model | Training<br>$F_{(1, 14)}$ = 10.64, ***p* = 0.0057** | Treatment<br>$F_{(1, 14)}$ = 12.69, ***p* = 0.0031** | Two-way interaction<br>$F_{(1, 14)}$ = 17.36, ***p* = 0.001** |
| Post-hoc Sidak's test | Sham<br>$t_{(14)}$ = 5.25, ***p* = 0.0002** | RSC lesion<br>$t_{(14)}$ = 0.64, *p* = 0.7814 | |
| **Panel F – Reward consumption** | | | |
| Sham<br>Two-way mixed-effects model | Training<br>$F_{(1, 7)}$ = 19.71, ***p* = 0.003** | Flavor<br>$F_{(1, 7)}$ = 4.05, *p* = 0.0841 | Two-way interaction<br>$F_{(1, 7)}$ = 6.12, ***p* = 0.0427** |
| Post-hoc Sidak's test | Initial Less Preferred<br>$t_{(7)}$ = 4.31, ***p* = 0.0071** | Initial Preferred<br>$t_{(7)}$ = 2.95, ***p* = 0.0424** | |
| RSC lesion<br>Two-way mixed-effects model | Training<br>$F_{(1, 7)}$ = 80.05, ***p* < 0.0001** | Flavor<br>$F_{(1, 7)}$ = 153.8, ***p* < 0.0001** | Two-way interaction<br>$F_{(1, 7)}$ = 74.52, ***p* < 0.0001** |
| Post-hoc Sidak's test | Initial Less Preferred<br>$t_{(7)}$ = 0.28, *p* = 0.9548 | Initial Preferred<br>$t_{(7)}$ = 10.63, ***p* < 0.0001** | |
| **Panel G – Initial Less Preferred correlation** | Sham<br>$r^2$ = 0.02, *p* = 0.74 | RSC lesion<br>$r^2$ = 0.36, *p* = 0.11 | |
| **Panel H – Initial Preferred correlation** | Sham<br>$r^2$ = 0.52, ***p* = 0.04** | RSC lesion<br>$r^2$ = 0.87, ***p* < 0.001** | |

| **Figure 7** | | | |
|---|---|---|---|
| Two-way mixed-effects model | Flavor<br>$F_{(1, 85)}$ = 3.89, *p* = 0.0517 | Change in preference<br>$F_{(2, 85)}$ = 5.78, ***p* = 0.0044** | Two-way interaction<br>$F_{(2, 85)}$ = 116.2, ***p* < 0.0001** |
| Post-hoc Sidak's test | Orange<br>$t_{(85)}$ = 10.61, ***p* < 0.0001** | Light Blue<br>$t_{(85)}$ = 0.95, *p* = 0.7211 | Dark Blue<br>$t_{(85)}$ = 10.94, ***p* < 0.0001** |
| **Panel B – Orange** | | | |
| One-sample t-test | Change in Initial Less Preferred consumption<br>$t_{(15)}$ = 0.69, *p* = 0.5007 | Change in Initial Preferred consumption<br>$t_{(15)}$ = 8.39, ***p* < 0.0001** | |
| **Panel C – Light Blue** | | | |
| One-sample t-test | Change in Initial Less Preferred consumption<br>$t_{(44)}$ = 88.88, ***p* < 0.0001** | Change in Initial Preferred consumption<br>$t_{(44)}$ = 5.47, ***p* < 0.0001** | |
| **Panel D – Dark Blue** | | | |

| | | |
|---|---|---|
| One-sample t-test | Change in Initial Less Preferred consumption<br>$t_{(26)} = 16.99$, ***p* < 0.0001** | Change in Initial Preferred consumption<br>$t_{(26)} = 2.23$, ***p* = 0.0348** |
